# A low-dimensional transcriptional code enables decoding of BMP/TGFβ signaling from single-cell transcriptomes

**DOI:** 10.64898/2026.04.28.721387

**Authors:** Guy Ilan, Inbal Eizenberg-Magar, Adi Wider, Jialong Jiang, Sisi Chen, Jong H. Park, Tsviya Olender, Matt Thomson, Yaron E. Antebi

**Author notes:** These authors contributed equally (Guy Ilan, Inbal Eizenberg-Magar).

## Abstract

Cells interpret complex extracellular environments through signaling pathways that compress diverse ligand inputs into a limited set of intracellular mediators. How this information is encoded in transcriptional responses, and whether it can be decoded to infer the original signaling environment, remains unclear. Here, we systematically map BMP/TGFβ transcriptional responses using single-cell RNA sequencing across 48 ligand-concentration conditions, generating a multi-ligand dose-response atlas. We find that, for each ligand, transcriptional responses collapse onto a one-dimensional, highly coordinated program in which concentration modulates response amplitude without altering gene identity. Across ligands, we find a small number of distinct, ligand-dependent transcriptional programs. We leverage this structured encoding to define a quantitative perception score that captures pathway activity at single-cell resolution. Finally, we train a machine learning model to decode extracellular ligand concentrations from single-cell transcriptomes. The model further generalizes to in vivo intestinal epithelium, where it reconstructs the spatial BMP gradient profile. Together, these results reveal a low-dimensional transcriptional code in BMP/TGFβ signaling that constrains cellular responses and enables quantitative inference of extracellular signaling environments from gene expression.

## Introduction

Cells continuously sense and respond to complex extracellular environments composed of multiple signaling ligands that vary in identity, concentration, and time. These signals are processed by intracellular signaling pathways that map diverse inputs onto a limited set of molecular mediators, forming a characteristic bottleneck between extracellular cues and transcriptional responses. How this compression shapes the encoding of environmental information, and to what extent transcriptional outputs retain sufficient information to recover the original signals, remain central open questions.

The Transforming Growth Factor-β (TGFβ) superfamily, which includes the TGFβ, BMP, and GDF ligands, provides a canonical example of such a system. More than 30 ligands signal through a shared set of receptors and converge on a small number of intracellular mediators, primarily SMAD1, 2, 3, 5, and 8, which regulate gene expression across diverse biological contexts [1–3]. Despite activating shared downstream mediators [4], individual ligands can elicit distinct cellular responses [5–7], raising the question of how specificity is encoded within a highly overlapping and constrained signaling network.

From an information-processing perspective, signaling pathways can be viewed as transformations between an input space, defined by extracellular ligand identities and concentrations, and an output space, defined by transcriptional states. A central challenge is to understand the structure of this mapping and determine whether high-dimensional extracellular environments are encoded in equally complex transcriptional responses, or whether the pathway imposes constraints that give rise to simpler, low-dimensional representations. While recent advances in single-cell sequencing have enabled detailed measurements of the output space, systematic and quantitative characterization of the input space remains limited. Resolving this relationship is essential for determining whether transcriptional measurements can be used to infer the signaling environment experienced by individual cells.

Here, we systematically map the transcriptional response of the BMP/TGFβ pathway using multiplexed single-cell RNA sequencing across 48 ligand conditions, generating a multi-ligand dose-response atlas. This dataset enables a quantitative characterization of the mapping between extracellular inputs and transcriptional outputs. We find that, for each ligand, transcriptional responses collapse onto a one-dimensional, highly coordinated program, in which concentration modulates response amplitude without altering gene identity and composition. Across ligands, these programs form a small number of distinct, ligand-specific transcriptional states.

Building on this structured representation, we define a quantitative measure of cellular signaling activity and develop a machine learning framework that decodes extracellular ligand concentrations from single-cell transcriptomes. We apply our decoding model to in vivo data from the intestinal villus to infer the spatial profile of a BMP signaling gradient. Together, our results reveal a low-dimensional transcriptional code for BMP/TGFβ signaling that constrains cellular responses and enables quantitative inference of extracellular signaling environments.

## Results

### Dose-dependent transcriptional responses to individual BMP/TGFβ ligands collapse onto one-dimensional trajectories

We first set out to systematically map the transcriptional response patterns generated by the BMP/TGFβ pathway. To this end, we used the epithelial NMuMG cell line, which exhibits robust responses to various signals from this pathway [1,8,9]. Cells were exposed to six ligands, BMP4, BMP6, BMP9, BMP10, GDF5, and TGFβ1, representing distinct structural and functional subgroups within the TGFβ/BMP pathway [10,11]. Given the morphogen-like behavior of BMP/TGFβ signaling, where distinct gene responses are expected across a concentration gradient [12,13], we stimulated cells with eight increasing concentrations spanning the dynamic range of each ligand (Table S1), yielding 48 distinct conditions (Figure 1A). We measured gene expression using high-throughput multiplexed single-cell RNA sequencing (MULTI-Seq) [14], at 3 and 6 hours following stimulation. Comparing the two timepoints, we observed highly similar transcriptional responses, indicating that the results are robust to the choice of timepoint within this window (Figure S1A-B). This analysis provides a comprehensive dataset of the transcriptional response profiles across ligand identities and concentrations.

**Figure 1.**
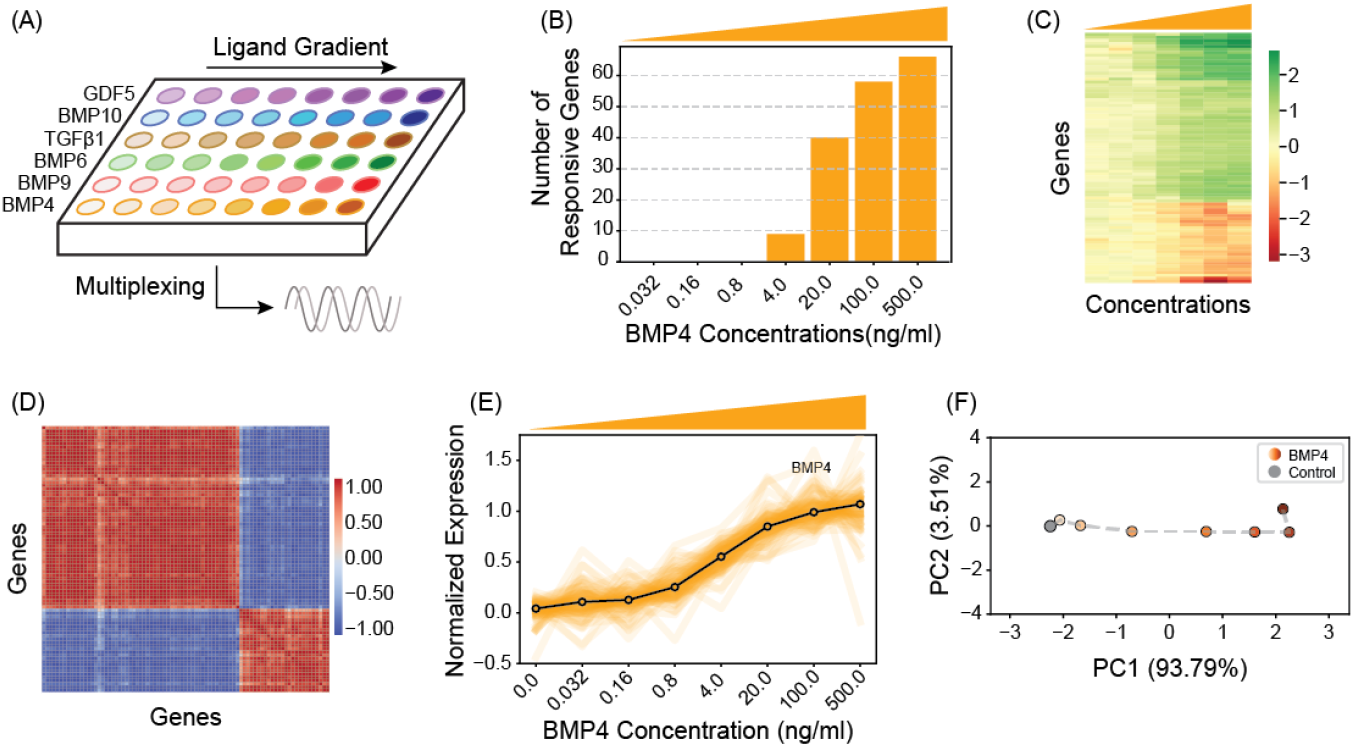
Dose-dependent transcriptional response to BMP4 defines a one-dimensional response trajectory. (A) Schematic representation of the experimental settings. NMuMG cells were exposed to six BMP/TGFβ ligands, each provided at eight increasing concentrations, depicted by a color gradient. Following 6 hours of ligand exposure, transcriptional responses were measured at the single-cell level using multiplexed RNA sequencing. (B) Number of significantly responsive genes identified at each BMP4 concentration (p-value < 0.05 & |Log_2_FC| > 1). (C) Heatmap display of log_2_ fold change in expression of 83 BMP4-responsive genes across the concentration gradient, with colors indicating upregulation (green) and downregulation (red). (D) Pearson correlation matrix of pairwise relationships between BMP4-responsive gene profiles across concentrations. Color indicates the direction of the correlation, positive (red) or negative (blue), and intensity represents the strength of the relationship. The high degree of correlation is consistent with a shared, unidirectional response structure. (E) Principal component analysis (PCA) of mean gene expression levels across BMP4 concentrations. Colored dots represent increasing ligand concentrations, ranging from low (light) to high (dark), connected by a dashed line to illustrate the concentration-dependent trajectory. A gray dot indicates the control sample (no added ligand). The first principal component (PC1) explains 93.79% of the variance and defines the dominant response direction. (F) Normalized mean expression profiles of individual BMP4-responsive genes across all tested concentrations, demonstrating collapse onto a shared sigmoidal dose-response trajectory. See also Figure S1.

We began by analyzing the response to BMP4, one of the most commonly studied BMP ligands with broad roles across biological contexts [15–19], focusing on the population-level response. For each concentration condition, we averaged the single-cell expression profiles to generate pseudo-bulk measurements. We identified responsive genes at each concentration using a significance threshold (p-value < 0.05) and an effect size cutoff (> 2-fold, see Methods). Examining the number of responding genes, we found it to increase with higher concentrations of BMP4 (Figure 1B). To capture the full response space, we defined the BMP4-responsive gene set as all genes that show a significant response in at least one concentration. Analyzing the genes across concentration increases statistical power and avoids imposing assumptions about gene monotonicity. In total, we identified 83 genes with robust responses to BMP4 and analyzed their pseudo-bulk response profile (Figure 1C).

Next, we characterized the full dose-response behavior of individual genes. Differences in gene profiles could arise from variations in sensitivities or more complex, non-monotonic responses. To assess this, we computed pairwise correlations between gene response profiles and performed hierarchical clustering on the resulting correlation matrix. All genes were either strongly positively or strongly negatively correlated (Figure 1D), indicating that all responsive genes share a single, universal response profile. Consistent with this, normalization of gene expression to their respective dynamic ranges revealed that all 83 BMP4-responsive genes collapse onto a shared sigmoidal curve (Figure 1E), exhibiting similar sensitivity to ligand concentration despite differences in the absolute expression levels. To further quantify this coordination, we performed principal component analysis (PCA) on the mean expression levels across concentrations (Figure 1F). The first principal component alone explained 93.7% of the variance, identifying a single dominant direction that captures the transcriptional response to BMP4. Together, these results demonstrate that BMP4 induces a coherent, effectively one-dimensional transcriptional program, in which ligand concentration modulates the amplitude of the response without altering its composition.

Extending our analysis to all ligands, we found that, in each case, responsive genes exhibited a coherent, graded, and co-linear response that collapsed onto a single trajectory, similar to BMP4 (Figure S1C-H). These results indicate that all tested BMP and TGFβ ligands induce a well-defined, effectively one-dimensional transcriptional program.

### BMP/TGFβ ligands induce specific transcriptional programs, resulting in distinct gene activation patterns

To compare the transcriptional programs elicited by all six tested BMP/TGFβ ligands, we analyzed the transcriptional response across the entire dataset. We identified all responsive genes for each ligand and combined them into a unified set of 733 genes. We then applied PCA to the pseudo-bulk expression levels of all responsive genes across all ligand-concentration conditions and visualized the data using the first three principal components, which together account for 95% (78.6%, 15%, 1.6%) of the total variance (Figure 2A). This analysis revealed that ligand responses occupy distinct regions of transcriptional space. In particular, BMP4, BMP6, and BMP9 share a common transcriptional program, whereas BMP10, TGFβ1, and GDF5 each form separate and distinct programs. Overall, the six ligands partitioned into four distinct transcriptional programs.

**Figure 2.**
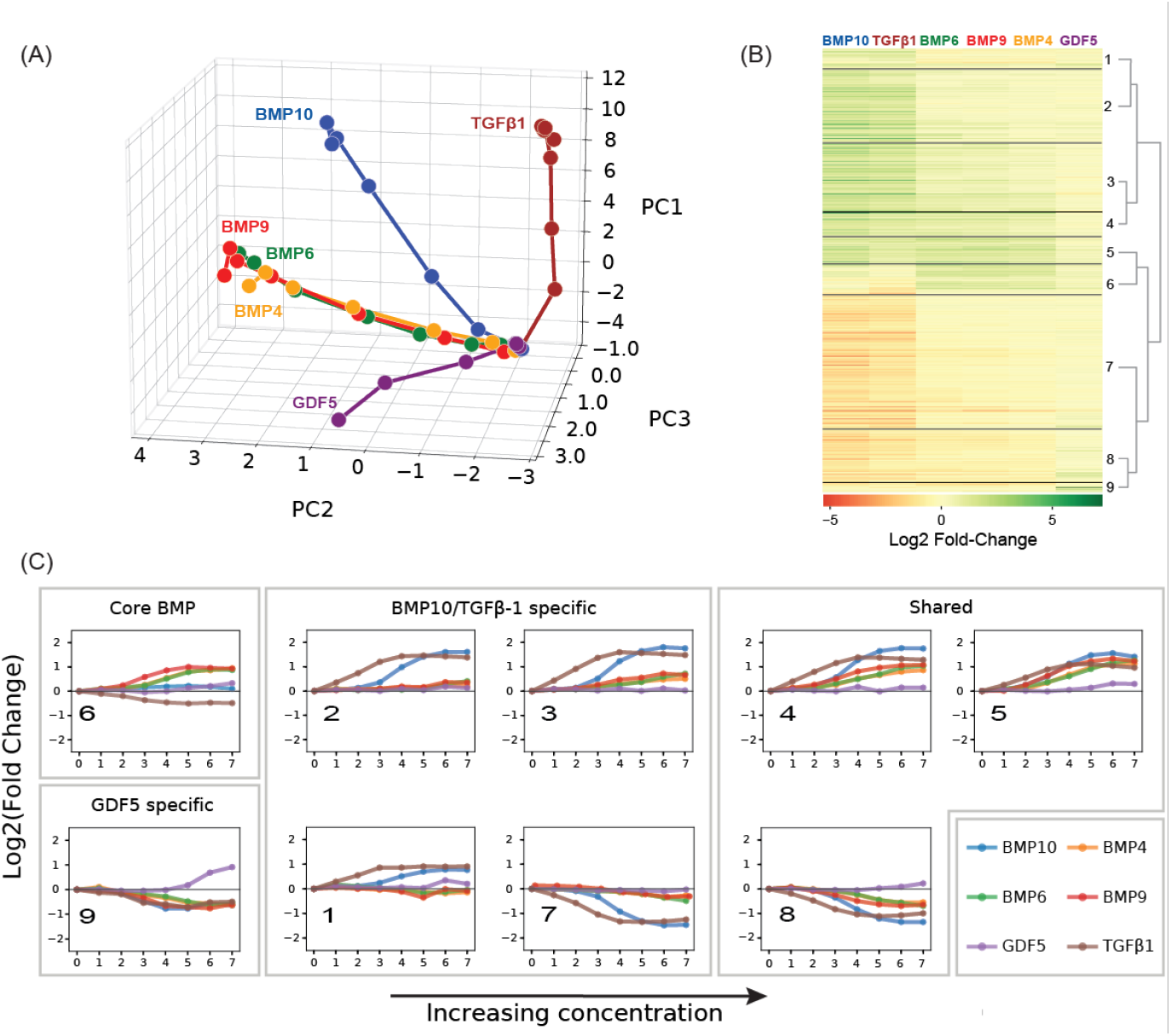
BMP/TGFβ ligands induce distinct transcriptional programs that shape gene activation patterns. (A) PCA of pseudo-bulk expression profiles across all ligand-concentration conditions. Each trajectory corresponds to a ligand, with dots representing increasing concentrations, interconnected to show concentration-dependence. Ligands induce transcriptional responses towards distinct directions in gene expression space, revealing separation into four primary response programs. (B) A hierarchically clustered heatmap showing the log_2_ fold change of 733 responsive genes (see Methods). Colors indicate upregulation (green) and downregulation (red) of varying intensities. The dendrogram shows clustering of genes into nine groups. (C) Gene clusters from panel B exhibit four response patterns: core BMP (BMP4, BMP6, and BMP9) responsive, BMP10/TGFβ1 responsive, GDF5 responsive, and globally responsive. Line plots show mean log_2_ fold change across increasing concentrations for each cluster, highlighting distinct activation dynamics. See also Figure S2.

To further characterize the gene-level structure underlying these programs, we analyzed the ligand-dependent response profiles of individual genes. For each gene, we fit its dose-response behavior using a Hill function and quantified its dynamic range as the fold-change between basal and maximal expression (Figure 2B). We then performed hierarchical clustering on these response profiles, identifying nine gene clusters that represent distinct patterns of ligand-specific activation (Figure 2C). These clusters group naturally into four classes that correspond to ligand-level programs. Cluster 6 contains genes that primarily respond to the core BMP ligands (BMP4, BMP6, and BMP9). Cluster 9 comprises a small set of genes uniquely responsive to GDF5. Clusters 1, 2, and 7 include genes that respond specifically to BMP10 and TGFβ1. Finally, clusters 3, 4, 5, and 8 contain genes that are broadly responsive across BMP and TGFβ ligands, representing a shared response program. The relationships between ligand programs are further supported by pairwise Pearson correlation analysis of ligand responses (Figure S2A). Together, these results demonstrate that BMP/TGFβ ligands organize gene expression into a small number of distinct gene clusters with specific ligand-dependent response patterns.

### Distinct gene activation patterns of ligands suggest functional diversity in the TGFβ superfamily

To interpret the functional consequences of the ligand-specific transcriptional programs, we performed Gene Ontology (GO) enrichment analysis on the gene clusters identified above, focusing on the four response patterns: BMP10/TGFβ1-specific, BMP-specific, GDF5-specific, and globally activated genes. This analysis revealed that while the TGFβ superfamily shares a core signaling architecture, individual ligands engage distinct transcriptional programs ranging from motility and tissue remodeling to homeostatic maintenance (Figure 3, S3).

**Figure 3.**
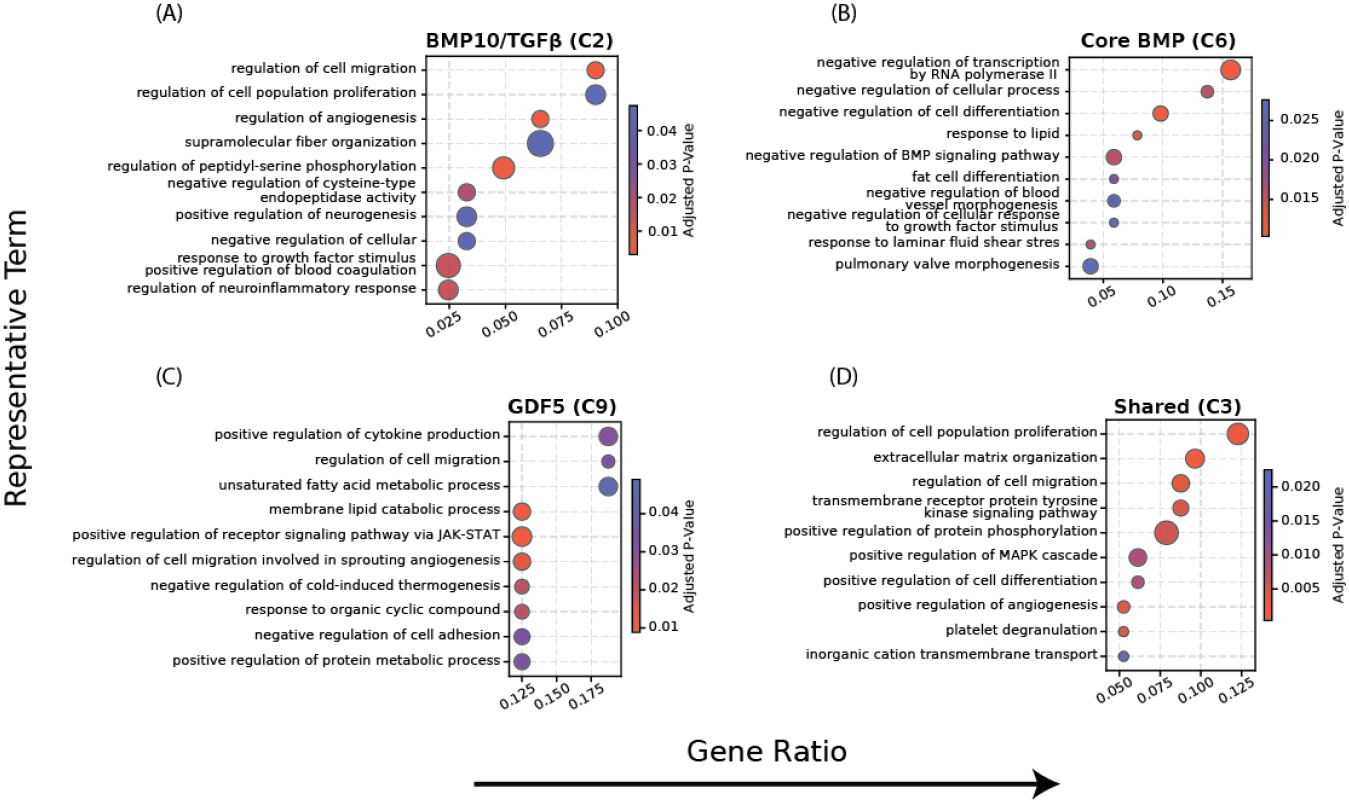
Gene Ontology analysis reveals functional programs underlying ligand-specific transcriptional responses. Representative GO terms for selected gene clusters corresponding to four transcriptional programs: BMP10/TGFβ1-specific (Cluster 2), core BMP (Cluster 6), GDF5-specific (Cluster 9), and shared responses (Cluster 3). Terms are grouped by similarity-based clustering and annotated by representative terms. Groups are ordered by gene ratio and colored by adjusted p-value. See also Figure S3.

The most significant transcriptional shift was observed in genes specifically responding to TGFβ1 and BMP10. These ligands drive a coordinated program that is characterized by the remodeling of both the cellular state and the extracellular environment, consistent with the acquisition of a migratory phenotype. The cells down-regulated genes involved in the targeted suppression of the homeostatic cell cycle and growth-factor responsiveness (CDK6, FGFR2), alongside the disassembly of established cytoskeletal organization (EPS8, EZR). In addition, we observed a significant decrease in transcripts associated with cellular connectivity and environmental sensing. This isolation is further reinforced by the suppression of guidance and contact-dependent sensing nodes (NRP1, EPHA4/EPHA7), supporting a loss of contact-mediated environmental awareness. In parallel, the cells upregulate genes required for active migration (SPHK1, NEDD9, RHOB) and the synthesis of a specialized extracellular scaffold (COL1A1, THBS1) required for invasiveness-associated matrix organization. The high enrichment of SERPINE1 and F3, alongside VEGFA, indicates a classic pro-angiogenic and hemostatic signature. Overall, these findings demonstrate that BMP10 and TGFβ1 induce a specific transition toward a migratory and invasive state, consistent with their established roles in epithelial-to-mesenchymal transition (EMT) and vascular remodeling [20–22].

In contrast to the invasive program driven by TGFβ1 and BMP10, GO analysis of genes specifically activated by core BMP ligands (BMP4, 6, and 9) revealed terms associated with the regulation of proliferation and differentiation. By upregulating potent transcriptional repressors of the ID family (ID1, ID2, ID3) and HEY1, these BMP ligands guide cellular differentiation and prevent the acquisition of undesired lineage identities. This reflects the known role of BMPs in maintaining cellular plasticity until the terminal stages of morphogenesis.

Our analysis further identified a distinct genetic program activated uniquely by GDF5. We found GDF5 enriched gene expression related to the regulation of various cytokines and growth factors, including VEGF, TGFβ, and IFNs, thereby priming the environment for angiogenesis and cytokine crosstalk. This corresponds to the known pro-angiogenic function of GDF5 [23–26].

Finally, we also identified globally responsive genes activated by both BMP and TGFβ ligands. This shared program suggests a sub-program that operates regardless of a specific ligand. In this group, we find a robust inhibitory feedback module that prevents runaway signaling and maintains homeostatic control. For example, the up-regulation of the intracellular inhibitor SMAD7 alongside extracellular antagonists NOG and NBL1, known sequesters of BMP ligands, indicates a conserved homeostatic mechanism designed to saturate the signaling response and prevent pathological over-stimulation of the TGFβ/BMP pathways. This internal regulation is coupled with a coordinated niche remodeling program that combines paracrine amplification, extracellular matrix (ECM) organization, and cell adhesion. The superfamily regulates matrix proteolysis and scaffold disassembly (CAPN2, TIMP1) and basement membrane-associated components (LAMB3, ITGB4) while upregulating potent mitogenic and pro-angiogenic growth factors (PDGFA/B, VEGFC, FGF18). Reflecting the high metabolic demand of this secretory state, we also observed the induction of the unfolded protein response (UPR) and ER stress markers (XBP1, EIF2AK3). Together, shared gene activation across the BMP/TGFβ superfamily enables cells to redefine their physical and signaling contexts within the tissue architecture.

### A single-cell perception score quantifies pathway activity from transcriptional data

Building on the observation that transcriptional responses are constrained along a single dominant direction for each ligand, we asked whether this structure could be used to quantify pathway activity at the single-cell level. For each ligand, we defined a response signature as the first principal component (PC1) derived from pseudo-bulk expression profiles across the concentration series. At the population level, this signature increases with ligand concentration, reflecting the graded nature of the response. We next projected individual single-cell transcriptomes onto the PC1-derived response signature, yielding a perception score that quantifies pathway activity in each cell (Figure S4A). As expected, cells exposed to higher ligand concentrations exhibit higher perception scores on average (Figure 4A), a relationship that was consistent across all six tested ligands (Figure S4B-F).

**Figure 4.**
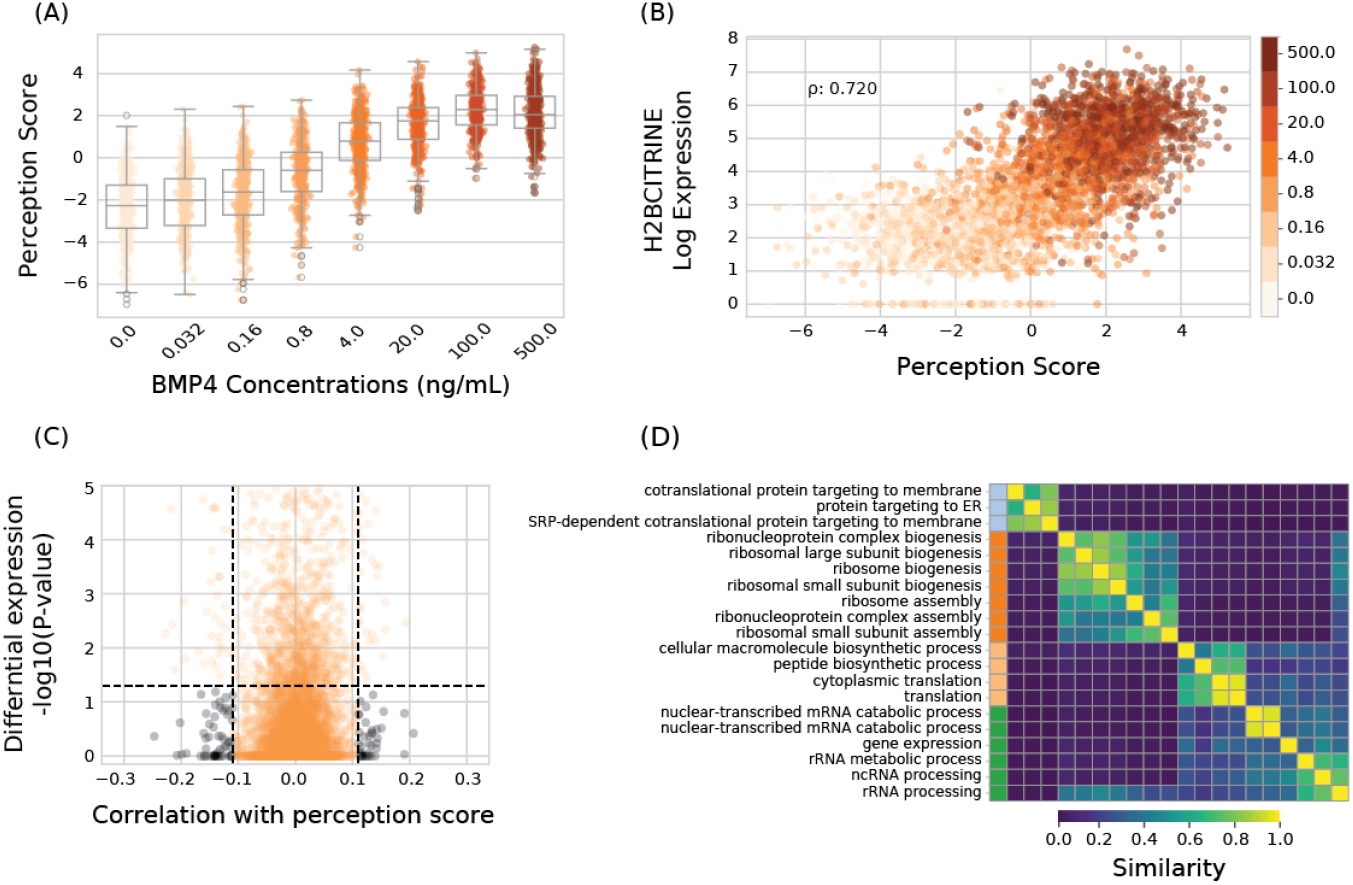
A single-cell perception score quantifies pathway activity and reveals cell-state-dependent variability. (A) Single-cell perception scores were calculated across BMP4 concentrations, showing a graded increase with ligand dose. (B) Perception score is plotted as a function of synthetic SMAD1/5/8-responsive reporter expression, demonstrating a strong correlation between the score and BMP pathway activity. (C) Partial correlation between gene expression and perception score (x-axis) and the p-value for differential expression (y-axis) are plotted for all non-responsive genes. Genes with significant correlation but no differential expression across conditions are highlighted, identifying candidate modulators of cell response. (D) Similarity clustering of significantly enriched Gene Ontology terms for the identified modulator genes, revealing enrichment for core cellular processes related to protein synthesis and RNA metabolism. See also Figure S4.

To validate the ability of our perception score to capture pathway activity, we used a synthetic SMAD1/5/8-responsive transcriptional reporter as an independent readout of pathway activation [9,27]. Reporter expression showed a strong correlation with the perception score across single cells (Figure 4B). This correlation remained significant within populations exposed to the same stimulus and was particularly pronounced at intermediate concentrations (Figure S4G), further demonstrating the capacity of the perception score to capture the signal response at the single-cell level.

While the extracellular signal is the primary determinant of the perception score, we observe substantial cell-to-cell heterogeneity even within populations exposed to the same stimulus. Notably, in our data, the most responsive cells at low stimulation overlapped with the least responsive cells at high stimulation. This suggests that, in addition to extracellular ligand concentration, intrinsic cellular factors contribute to the observed variability in signaling response. To further investigate the intercellular variability and identify the contributing factors, we identified all genes whose expression is correlated with the perception score after controlling for ligand concentration. This set of genes could include both responsive genes that increase with pathway activity and modulating genes that control the way cells process and respond to signals. To focus on potential modulators rather than responsive genes, we excluded genes that exhibited any dependence on stimulus level (Figure 4C). From this analysis, we identified a set of 100 possible modulating genes. Performing GO analysis on these genes, we identified consistent enrichment across ligands for fundamental processes, including ribosomal biogenesis, RNA processing, and protein transportation (Figure 4D). These results suggest that the basal cellular state, particularly processes related to biosynthetic capacity, influences how individual cells interpret and respond to extracellular signals.

### A machine learning framework enables the reconstruction of extracellular ligand concentrations from single-cell transcriptomes

The perception score provides a quantitative measure of pathway activity at the single-cell level. However, this score reflects a cell-specific response that depends not only on extracellular ligand concentration but also on intrinsic cellular context, including expression of receptor, signaling components, and cell-state variables. Such cellular context genes may not respond to a stimulus, but rather modulate the response, resulting in variable perception scores in cells exposed to identical stimuli. We therefore asked whether incorporating both responsive and modulating genes could enable direct inference of the extracellular signal and reduce the observed variability.

We trained a machine learning (ML) model to predict ligand concentration from single-cell transcriptomes. We selected the top 2,250 highly variable genes (HVGs), capturing both responsive and non-responsive genes, and trained a gradient-boosted decision tree model (XGBoost) to classify cells into eight concentration classes. Training was performed on a combined set of cells exposed to BMP4, BMP6, and BMP9, which share a common response program (Figure S5A). Model performance was evaluated on a held-out test set using cumulative Receiver Operating Characteristic (ROC) curve analysis and the confusion matrix (Figure 5A, S5B-C). The model demonstrates robust performance across all concentrations, achieving an area under the curve (AUC) exceeding 0.924, which indicates high predictive accuracy of the model.

**Figure 5.**
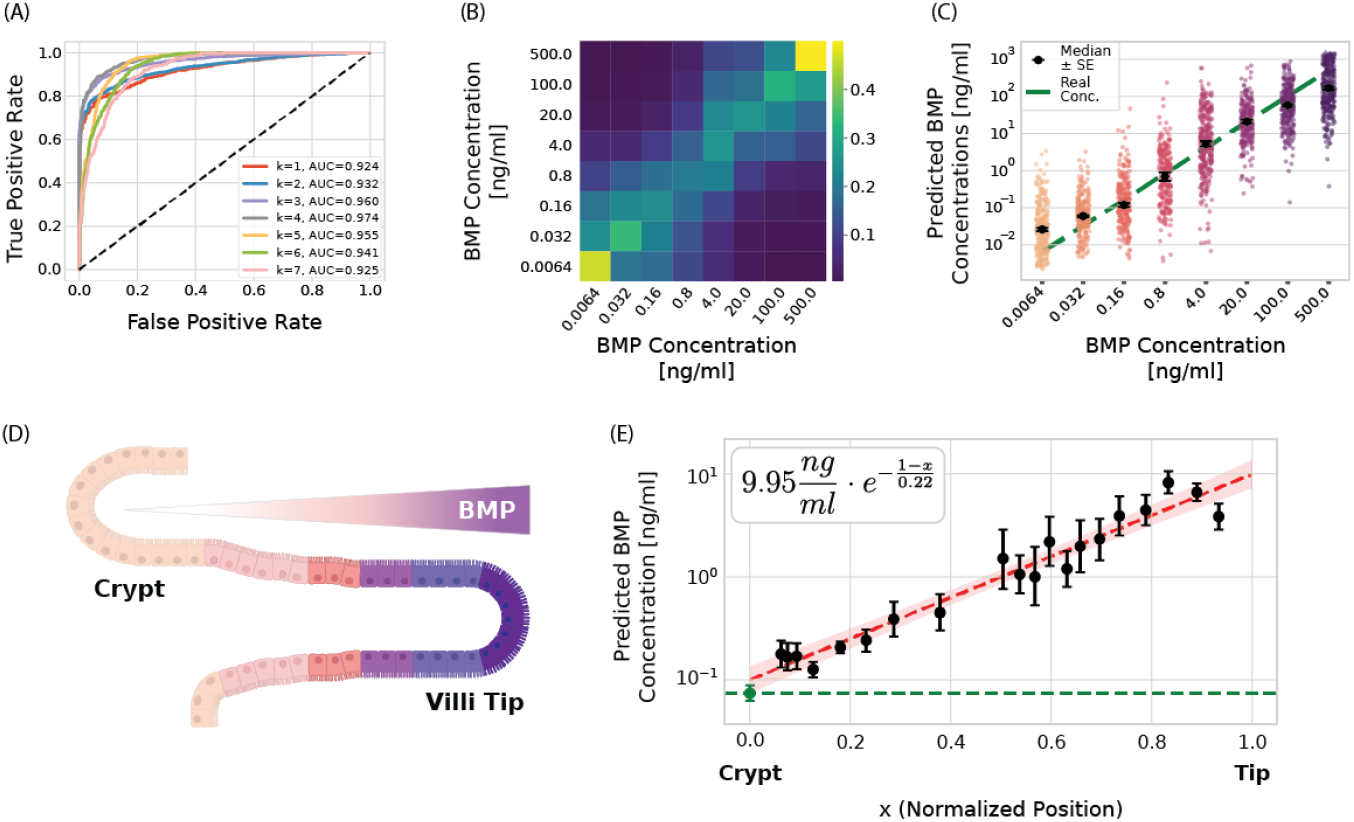
Machine learning enables reconstruction of extracellular ligand concentrations from single-cell transcriptomes. (A) Cumulative Receiver Operating Characteristic (ROC) curve analysis for classification of concentration classes, demonstrating robust predictive performance across all concentrations (AUC > 0.924). (B) Aggregated predicted class probabilities for cells exposed to each concentration. Probability mass shifts toward higher concentration classes as the true ligand concentration increases, with the highest confidence at the lowest and highest concentrations. (C) Continuous predicted BMP concentrations for individual cells, derived from probabilistic model outputs, showing accurate recovery of input concentrations at single-cell resolution. (D) Schematic of the intestinal crypt-villus axis, where a spatial gradient of BMP signals regulates the epithelial villi cell differentiation. (E) Predicted BMP concentrations mapped along the crypt-villus axis in vivo. Median predicted concentrations (± s.e.) are shown for spatially binned cells. Green dashed line represents the crypt; red dashed line represents the linear fit between the variables. The reconstructed gradient follows an exponential profile (R^2^ = 0.95), enabling quantitative estimation of effective ligand levels and gradient length scale. See also Figure S5.

To further characterize model predictions, we examined the full set of predicted probabilities over concentration classes for each cell. For each concentration, we then aggregated all cells exposed to that condition and calculated the distribution of predicted probabilities for each concentration class. These distributions generally follow the true ligand concentration increase, with the most confident predictions occurring at the extreme lowest and highest concentrations (Figure 5B). We also find that predictions are centered around the true concentration value, with large deviations being relatively rare. We leveraged this probabilistic structure to derive a continuous prediction of ligand level for each cell. We fitted the predicted concentration-class probabilities for each cell to a Gaussian distribution and extracted its mean to infer a continuous concentration. This approach enabled accurate reconstruction of ligand concentration at single-cell resolution (Figure 5C).

Notably, and in contrast to the perception score defined earlier, predicted concentrations exhibited reduced variability across cells exposed to the same condition. Furthermore, no overlap was found between the distributions of maximal and minimal activation, demonstrating the model’s ability to fully distinguish between treated and untreated cells. This indicates that the ML model captures additional information beyond the primary response direction, effectively integrating cellular context to improve inference of extracellular signals. To quantify this reduction in variability, we compared the standardized variance of the perception score with that of the predicted concentration across conditions. (Figure S5D). Indeed, predicted concentrations consistently showed lower variance, supporting improved separation of cellular states.

### Applying the ligand inference model on an in vivo intestinal transcriptome data quantitatively recovers the BMP gradient

To assess whether this framework generalizes beyond controlled in vitro conditions, we applied the trained model to an independent single-cell dataset of intestinal epithelial cells, where a spatial BMP gradient is known to exist along the crypt-villus axis (Figure 5D). This system is particularly suited to our analysis for several reasons. First, it is an important and well-studied biological system for which high-quality single-cell RNA-seq datasets are available. Second, most variations occur along a single direction, from the bottom of the villus to its tip, resulting in an effectively one-dimensional system. Third, the BMP gradient is well known, with high levels at the tip and low levels at the bottom [28,29]. However, while some work has attempted to determine the gradient using physical models [30], the shape of the gradient is still not fully understood. Fourth, it is a temporally stable developmental system, with a stable BMP gradient that keeps its shape and structure. Thus, the intestinal villi provide an opportunity to test whether our model can recapitulate the direction of an in vivo BMP gradient and generate predictions about its shape.

We applied our pre-trained model on a single-cell intestinal epithelium dataset [31] and extracted a predicted BMP value for each single cell. Using a previously established zonation metric to infer spatial position [32], we mapped predicted ligand exposure along the crypt-villus axis. We then binned all cells based on spatial proximity into 20 bins, each containing 57 cells, and estimated the median predicted concentration for each bin. The model revealed a smooth gradient of BMP concentrations increasing towards the villus tip, consistent with current knowledge [28,29]. Fitting the reconstructed profile to an exponential decay model yielded a strong fit (*R* _*2*_*=0*.95), consistent with diffusion-like gradient formation. This analysis provided several quantitative estimates. We find an effective BMP ligand concentration of approximately 9.95 ng/ml at the villus tip (95% confidence interval of [7.35ng/ml, 13.48ng/ml]). Additionally, we can estimate the gradient lengthscale to be 22% of the villus length. Using an average villus length of about 250um [33], this gives a characteristic lengthscale of 55μm. Overall, these results further demonstrate that transcriptional data can be used to reconstruct extracellular signaling environments and infer the spatial structure of signaling gradients.

## Discussion

Cells are continuously exposed to complex extracellular environments composed of multiple signaling ligands across a range of concentrations. Understanding how cells encode these signals into transcriptional responses, and whether this encoding preserves sufficient information to induce ligand-specific responses, remains a central challenge. In this work, we systematically characterized transcriptional responses to various BMP/TGFβ ligands across their dynamic response ranges, generating a quantitative single-cell atlas of pathway activity. This dataset provides a controlled framework for linking extracellular inputs to transcriptional outputs. Our work highlights the value of generating additional systematic datasets across signaling pathways, cell types, and cellular contexts for dissecting cell-cell communication.

Analyzing our data reveals a distinct genetic signature for individual ligands, with highly constrained transcriptional responses within each signature. Increasing ligand concentration does not alter the composition of responding genes, but instead scales their coordinated expression along a shared response trajectory, resulting in a correlated increase in the response. Across ligands, these responses organize into a small number of distinct transcriptional programs, with BMP4, BMP6, and BMP9 defining a shared core BMP program, and BMP10, TGFβ1, and GDF5 defining separate programs. This organization demonstrates that BMP/TGFβ signaling encodes diverse extracellular inputs within a limited set of transcriptional states, enabling discrimination between ligand classes despite convergence onto shared intracellular mediators. While this framework allows classification of ligands based on transcriptional responses, an open question remains how individual ligand identities are resolved within this constrained representation.

A notable outcome of this analysis is that BMP10 defines a distinct transcriptional trajectory that lies between the core BMP and TGFβ1 response directions. Rather than clustering with either group, BMP10 occupies an intermediate position in transcriptional space, suggesting that it engages a partially overlapping but distinct signaling program. This positioning suggests that BMP10 may represent an intermediate functional state between canonical

BMP and TGFβ signaling. Whether this signature reflects an integrated state combining features of both pathways or a distinct transcriptional program requires further investigation. More broadly, the intermediate positioning of BMP10 raises the possibility that similar hybrid transcriptional states could be generated through combinatorial BMP and TGFβ ligand exposure, providing a potential mechanism for expanding signaling diversity within a constrained pathway architecture.

Building on the identification of ligand-dependent transcriptional programs, we leveraged the low-dimensional structure of the response to define a perception score that quantifies pathway activity at single-cell resolution. This score enables direct measurement of BMP/TGFβ pathway activity from transcriptomic data without the need for engineered reporters. Applying this framework revealed substantial heterogeneity in signaling responses among cells exposed to the same extracellular conditions. This variability reflects differences in basal cellular state, as genes associated with biosynthetic capacity, RNA processing, and proteostasis correlate with perception score independently of ligand concentration. These results indicate that the intrinsic cellular state is a key determinant of how cells interpret and respond to extracellular signals. Together, our findings link cellular pathway perception, ligand-responsive genes, and context-modulating genes, defining the basis by which transcriptional responses can reflect the underlying signaling environment.

Our findings demonstrate that this structured transcriptional response can be used to infer the underlying extracellular signaling environment. By integrating both responsive and context-modulating genes, we developed a machine learning framework that infers extracellular ligand levels from single-cell transcriptomes. This approach extends the perception score by incorporating intrinsic cellular variability, enabling more accurate inference of ligand dose at single-cell resolution. To assess generalizability beyond the in vitro system, we applied the model to an independent in vivo dataset of mouse intestinal enterocytes. While BMP gradients across the intestinal villus are well established, their quantitative spatial profile at single-cell resolution remains poorly defined. When applied to intestinal epithelial cells, the model recapitulates the spatial BMP gradient along the crypt-villus axis and reveals an exponential relationship between cellular position and predicted ligand exposure. This corresponds to an inferred BMP concentration at the villus tip of approximately 9.95 ng/ml with a gradient lengthscale of ~55 μm. Thus, our model provides a quantitative estimate of BMP exposure in vivo, derived from transcriptional response similarity, rather than a direct biochemical measurement. Together, these results demonstrate that models trained on controlled in vitro perturbations can recover physiologically relevant signaling gradients providing, for the first time, an estimate of the effective concentration of BMP ligands in an in vivo setting.

In summary, our work establishes a framework for decoding extracellular signaling environments from transcriptional data by leveraging the low-dimensional structure of pathway responses. By integrating both responsive and context-modulating gene sets, we demonstrate that machine learning models trained on controlled perturbations can quantitatively reconstruct extracellular ligand concentrations and further generalize to infer physiological signaling gradients in vivo. More broadly, these findings demonstrate that signaling pathways encode environmental information in structured transcriptional responses that can be quantitatively decoded to reconstruct the underlying signaling environment. Expanding this approach across additional pathways, cell types, and temporal regimes will enable the characterization of increasingly complex signaling environments. As more high-resolution datasets become available, more advanced modeling approaches, including generative frameworks, may enable synthetic reconstruction of entire signaling landscapes. Ultimately, this methodology provides a quantitative framework for uncovering the principles by which cells interpret and integrate extracellular cues across development, tissue homeostasis, and disease.

## Acknowledgments

This work was done with critical advice and help from the AI Hub at the Weizmann Institute of Science. YEA is supported by a research grant from the Sygnet Fund. This work was also supported by the Schwartz Reisman Collaborative Science Program, which is supported by the Gerald Schwartz and Heather Reisman Foundation.

## Methods

### Cell line and tissue culture

NMuMG (NAMRU Mouse Mammary Gland cells, female) (CRL-1636) were purchased from ATCC (CRL-1636). The cells were transfected with an H2B-mCitrine fluorescent reporter driven by BMP Responsive Element (BRE) [9,27]. Cells were grown in a humidity-controlled chamber at 37°C with 5% CO2 and cultured in DMEM supplemented with 10% FBS, 1 mM sodium pyruvate, 100 unit/ml penicillin, 100 μg/ml streptomycin, 2 mM L-glutamine, and 1x MEM non-essential amino acids.

For single-cell RNA sequencing (scRNA-seq), reporter NMuMG cells were cultured at 40% confluency in 96-well plates. Following 24h, cells were washed with phosphate-buffered saline (PBS) and added with fresh media containing ligands at varying concentrations, spanning the entire dynamic range of each ligand (Table S1). Specifically, cells were exposed to six different ligands, each administered at eight different concentrations, resulting in a total of 48 experimental conditions. All recombinant ligands were acquired from R&D Systems (Table S2).

### Single-cell RNA sample preparation, library construction, and sequencing

Library construction and the resulting scRNA-seq data were generated through a collaboration with the Thomson lab at Caltech. Briefly, reporter NMuMG cells were plated at 40% confluency and cultured for 24h. After 24h, the cells were added with ligands as indicated and cultured for an additional 6 hours. Cells were further dissociated into a single-cell suspension using TrypLE (3 min at 37°C), washed, and resuspended in PBS. To achieve high-throughput multiplexed sequencing, we utilized the MULTI-seq method [14]. For sample multiplexing, cells were incubated with barcoding solution containing equimolar amounts of LMO anchor and sample barcode oligonucleotides for 5 min at 4°C. Samples were further added with LMO co-anchor for 5 min at 4°C, followed by the addition of 10% BSA to quench the LMO reaction. Cells were washed, resuspended in PBS containing 1% BSA, and pooled. Finally, cells were filtered, counted, adjusted to 1×10^6^ cells/ml, and loaded on 10x Genomics lanes. Sequencing libraries were constructed using the 10x Genomics

Single Cell v3.1 3’ platform and sequenced on the Illumina NovaSeq 6000 platform. Reads are aligned to the mouse mm10 transcriptome with Cell Ranger v5.0.1 (10x Genomics). We used a custom Python pipeline to assign cells to each condition based on the sequenced tags, following the pipeline of MULTI-seq [14].

### Single-cell RNA-sequencing analysis

The resulting single-cell RNA-seq data contained approx. 28k cells and 28k detected genes. As a quality control, we excluded cells with fewer than 4000 detected genes (genes with UMI count > 0) and mitochondrial reads percent per cell higher than 10%. To account for the variation in the number of transcripts captured per cell, unique molecular identifier (UMI) counts were log2-normalized using this equation:

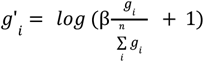

where *g*_*i*_ stands for the number of transcripts per i-th gene, n being the total number of analyzed genes and scaling factor-β stands for the mean of total transcripts per cell. After data processing, we remain with ~20k high-quality cells for further analysis.

### Identification of responsive genes for BMP ligands

We identified differentially expressed genes (DEG) in response to the six distinct ligands (BMP10, TGFβ-1, BMP4, GDF5, BMP6, and BMP9) across varying concentrations by comparing against the control conditions. Initially, we used the Kolmogorov-Smirnov Test (KS-test), a statistical method that enables a comparison of gene expression distributions by testing the distribution shift from the control. Notably, for each of the six ligands, we analyzed eight increasing concentrations in a gradient pattern, resulting in a total of 48 conditions. These conditions include a concentration of 0, where cells were not stimulated, giving rise to six control conditions. To identify genes responding to ligand treatment, we performed a two-sample KS test for each of the 42 experimental conditions (representing all ligand-concentration combinations excluding the six control replicates) against the pooled control distribution. Prior to testing, the dataset was filtered to include only genes with non-zero expression across the sample population, resulting in a candidate set of 18,000 features. This pre-filtering step was implemented to reduce the multiple testing burden and enhance statistical power. Resulting p-values were adjusted for multiple comparisons using the Benjamini-Hochberg False Discovery Rate (FDR) procedure. Genes with an adjusted

p-value < 0.05 (6,249 genes) were considered statistically significant. To further characterize the directionality and magnitude of these responses, we calculated the log_2_ fold-change of the mean expression for each significant gene in every treatment condition relative to the mean of the pooled control samples. Ultimately, DEG for each ligand was defined as those exhibiting both a statistically significant difference and a minimum two-fold change in expression in at least one of the ligand’s concentrations. This process generated sets of responsive genes, including both upregulated and downregulated genes for each ligand. These refined gene sets served as the foundation for our downstream analyses.

### Four-parameter logistic curve derived log_2_ fold-change values

To derive ligand-specific scalar fold-change values for each gene, we first established a global response model using the mean expression-derived Principal Component 1 (PC1) for each ligand (Figure 1E). This PC1 profile was fitted to a four-parameter logistic (4PL) curve to determine shared *EC50*and Hill slope (*n*) parameters. Then, pseudo-bulk expression profiles for 733 DEG’s were computed per ligand. To estimate gene-specific response magnitudes, each gene’s profile was min-max scaled, and a 4PL model was applied with the *EC50*and *n fix*ed to the previously determined PC1 values. The resulting fitted plateaus were rescaled to their original units, and final fold-changes were calculated from these plateaus according to this equation:

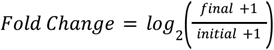

Where *final is* the fitted high concentration plateau value and *initial is* the fitted low concentration plateau value for each gene.

### Normalized gene expression levels

To compare distinct genes with different activation levels, we calculated the normalized expression levels of all DEG’s, defined as the expression levels relative to the full response range. The initial/final plateaus calculated for each gene by the 4PL fit were used in order to scale the values of all genes between their respective fitted range:

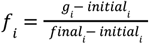

where *g* _*i*_ is the pseudo-bulk expression value of gene *i*. This formula adjusts *g* _*i*_ *val*ue by subtracting the basal level (*initial*_*i*_) and then scaling the result to the range defined by the saturation level (*final* _*i*_) minus the basal level (*initial* _*i*_). The normalized expression value thereby represents the position of the gene’s expression level on its response curve, ranging from 0 (at the basal level) to 1 (at the maximum level), thus providing a comparative measure of gene activity. We note that the normalized expression level is calculated equivalently to upregulated and downregulated responsive genes.

### Clustering of differentially expressed genes

To identify distinct gene response patterns, the gene-by-ligand fold-change matrix was Z-score standardized on a per-gene basis. This transformation ensures that the subsequent clustering is driven by relative response dynamics across ligands rather than the absolute magnitude of expression. We then performed hierarchical clustering using Ward’s linkage and Euclidean distance, with optimal leaf ordering applied to both genes and ligands to refine the visualization.

### Enrichment analysis

The nine gene clusters identified in the initial clustering analysis (Figure 2) served as the input gene sets for Gene Ontology (GO) enrichment and subsequent term aggregation. Enrichment analysis against the GO Biological Process (2021) database was performed for each cluster using *gseapy* [34], with the background set to all non-zero genes in the original dataset. Statistical significance was defined using an FDR threshold of *P < 0… 05*, derived from a hypergeometric test. In order to reduce redundancy and enhance the interpretability of the results, pairwise semantic similarities between significant GO terms were calculated in R using the Wang measure in the *GOSemSim* package [35]. The resulting similarity matrices were processed through a custom Python pipeline that converted semantic similarity (*S*) into a distance metric (1 − *S*) for hierarchical clustering via Ward’s linkage. To determine the optimal number of groups (*k*) for each cluster, we implemented a dynamic search that identified the highest cluster resolution that satisfied a zero-singleton constraint. This ensured that every reported biological theme represented a community of functionally analogous terms rather than isolated results. Each final group was assigned a representative term based on a multi-tier priority sort: highest Gene Ratio (the proportion of the input cluster mapped to the term), followed by the Combined Score, lowest adjusted p-value, and GO hierarchy depth. To link these abstract functional themes back to the molecular data, we identified the “top genes” for each group by calculating the most frequently occurring gene symbols across all GO terms in each group. Files with group summaries included representative labels, term counts, and driving genetic components were exported for downstream biological interpretation.

### Perception score via single cell projection on PCA space

The projection of single-cell data onto the principal component analysis (PCA) space was executed in two stages. Initially, the principal component model was fitted utilizing the log_2_-transformed mean expression values of the ligand-responsive genes for each of the eight concentration levels (seven concentrations plus a control). Then, log_2_-transformed single-cell expression data were projected onto that same PCA model to get per-cell PC embeddings. The cells’ respective PC1 values were designated as their Perception Score, which serves to estimate the cell’s ligand activation state relative to the complete spectrum of observed responses.

### Synthetic marker gene - H2BCITRINE

The NMuMG cell line used in this study contains a synthetic transcriptional reporter driven by a promoter with multiple SMAD1/5/8-responsive elements, which serve as downstream effectors of BMP signaling. Activation of this reporter results in the expression of a Citrine fluorescent protein, providing a transcriptional readout of BMP pathway activity. The expression values of the reporter gene (“H2BCITRINE”) were normalized together with the rest of the transcriptomic data. To avoid bias in downstream analyses, particularly in the construction of predictive models, this gene was excluded from all modeling steps, ensuring that the inference of pathway activity was based solely on endogenous gene expression.

### Analysis of unresponsive modulators of signaling

We restricted our analysis to expressed genes (>0 counts) to identify treatment-unresponsive modulators of signaling perception. Partial correlations were calculated between gene expression and cellular perception scores, controlling for ligand concentration by correlating the residuals of each measure with the one-hot encoded categorial concentration variable. To isolate unresponsive features, we compared treated versus control distributions using a KS test. We selected the top 100 genes per ligand exhibiting high perception correlation but no significant distributional deviation. Gene Ontology (GO) enrichment was performed on these subsets, and terms conserved across all ligands were analyzed for pairwise semantic similarity and clustered to visualize functional groupings.

## Data preprocessing and quality control before model training

To ensure a robust training set for the machine learning model, we performed stringent quality control and normalization on the single-cell RNA sequencing data. We first filtered cells based on viability and sequencing depth, retaining only those with fewer than 10% mitochondrial reads and a minimum of 4,000 detected genes. Samples with unknown labels were removed, and potential gene-naming ambiguities were resolved by identifying duplicate gene entries and retaining only the entry with the highest total UMI count.

## Normalization and feature selection

Following cell-level filtering, we performed total count normalization to a target sum of 10,000 reads per cell, followed by a *log*_2_ (x + 1) transformation. To focus the model on biologically informative signals, we restricted the feature space to 2,250 highly variable genes identified via the Seurat v3 [36].

### Training data preparation

The continuous ligand concentrations were transformed into eight ordinal ranks (*C*_0_,…,*C*_7_) to serve as classification targets. The processed dataset was then split into training (80%) and testing (20%) sets using a stratified approach to maintain consistent concentration distributions across both subsets.

### XGBoost model training and hyperparameter optimization

We trained an XGBoost multiclass model to classify single-cell transcriptomic profiles (2,250 gene features per cell) into eight concentration levels (1-8). Hyperparameters were optimized with Optuna using a Tree-structured Parzen Estimator (TPE) sampler, a Bayesian strategy that proposes promising hyperparameter values based on prior trial outcomes, and a median pruner, which stops underperforming trials early by comparing intermediate results to the running median of completed trials. The objective minimized 5-fold cross-validated multiclass log-loss (mlogloss) with early stopping (20 rounds). Tuning was performed stepwise across four parameter groups: tree complexity (max_depth, min_child_weight), subsampling (subsample, colsample_bytree), regularization (alpha, lambda, gamma), and boosting dynamics (learning_rate, num_boost_round), carrying forward best values between groups. The final model was retrained on the full training set using the best hyperparameters found through tuning.

### Receiver Operating Characteristic (ROC) curve analysis

To evaluate the performance of the XGBoost model across the ordinal concentration gradient, we implemented a cumulative Receiver Operating Characteristic (ROC) analysis. Given the ordinal nature of the eight concentration classes *(C*_*0*_,…, *C*_*7*_ ), we assessed the model’s ability to distinguish between concentrations at increasing thresholds. For each threshold *k* ∈ {1,…, 7} binary labels were generated by partitioning the data into two groups: cells exposed to concentrations *Y ≥ C* _*k*_ (positive class) and those exposed to *Y < C*_*k*_ (negative class). The cumulative probability for each threshold was calculated by summing the predicted class probabilities from the XGBoost output: 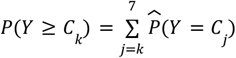 curves were then constructed by plotting the True Positive Rate (TPR) against the False Positive Rate (FPR) for each cumulative threshold, and the Area Under the Curve (AUC) was calculated to quantify the classification accuracy across the concentration spectrum.

### Aggregated probability predictions

To assess classification confidence across the ordinal scale, we grouped single-cell samples by their ground-truth exposure concentrations *(C*_*0*_,…, *C*_*7*_). For each group, we calculated the mean predicted probability for all eight potential classes. This aggregation resulted in an 8 × 8 matrix, where each row *i* represents the average softmax output vector for all cells truly belonging to concentration neighboring concentration levels. *C* _*i*_, revealing the distribution of probability mass across

### Concentration inference and predicted-concentration calculation

To refine the discrete output of the XGBoost model into a continuous exposure metric and mitigate boundary effects at the extreme ends of the concentration gradient, we implemented a regularized Gaussian reconstruction. We assumed that the predicted class probabilities *p* for each cell follow a discretized normal distribution across the concentration indices *i ∈ {0,…, 7}*. A global dispersion parameter (σ) was first estimated by averaging the standard deviations of Gaussian fits from cells with peak probabilities in the central classes (_2_ *≤ peak ≤ 5)*.

For each cell, we performed a regularized least-squares optimization to find the optimal distribution mean (μ). The objective function minimized the residual between the predicted probabilities and the binned Gaussian cumulative distribution function (CDF), defined as:

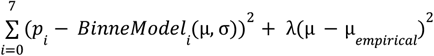

where *BinnedModel* _*i*_ is the probability mass within bin *i* calculated from the Gaussian CDF, μ _*empirical*_ *= ∑ i · p* _*i*_ is the empirical center of mass, and λ represents the regularization strength (0.075). The resulting fitted mean was then transformed into the physical concentration scale using the linear relationship of the experimental design:

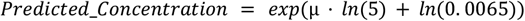

This approach allowed for the interpolation of predicted-concentrations beyond the discrete class centers. Next, we aggregated these predictions across experimental groups, and calculated group-level medians of the log-transformed values. Standard errors (SE) for these medians were estimated using a bootstrap approach (*n* = 1000 iterations). These SE values were then back-transformed to the original scale to provide upper and lower error bounds for the predicted-concentration estimates.

### Reconstruction of gut villus BMP gradient

Raw UMI counts of 1383 enterocyte and crypt cells were filtered and spatially reconstructed by Moor Et al. [32], single-cell spatial coordinates were assigned using their reconstruction algorithm. The spatial UMI data underwent the same normalization and scaling as the training data to ensure model compatibility. We then applied the trained model to these spatial profiles to predict exposure probabilities, followed by the previously described predicted-concentration calculation and regularized Gaussian reconstruction.

The original continuous spatial coordinates were transformed via linear interpolation between defined zone cutoffs to calculate a “continuous zone” score on the scale between 0-6 and divided by 6 to get a “normalized position” score between 0-1. Cells identified as belonging to the crypt (Zone 0) were assigned a fixed coordinate of 0, while all other cells were partitioned into 20 equal-sized bins using a quantile-based approach. For each spatial bin and the crypt zone, we calculated the median reconstructed predicted-concentration and estimated standard errors as previously described.

### Spatial gradient modeling and decay analysis

To quantify the concentration gradient along the villus axis, we modeled the relationship between the normalized spatial coordinates *(x)* and the reconstructed predicted-concentrations *(C)* as an exponential decay process, *C(x) = C*_*source*_ *· e*^*−k(*1−*x)*^. We performed a linear regression on the log-transformed median pseudo-concentrations of the spatial bins:

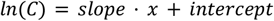

where the decay constant, *k*, and concentration at the source boundary *C* _*source*_ *ca*n be estimated using:

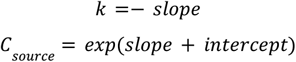

To account for uncertainty in this estimate, we calculated a 95% confidence interval by applying the residual standard error (RSE) to the prediction in log-space before back-transformation. The standard error of the prediction at *(x = 1… 0)* was determined by:

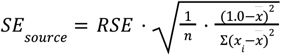

We further characterized the profile using the characteristic lengthscale (λ), which represents the distance from the source at which the concentration decreases to 1/*e(*≈ *37%) of* its initial value. This lengthscale was derived as the reciprocal of the absolute slope (the decay constant, *k*), λ *= 1*/*k*.

## Supplementary Figures

**Figure S1.**
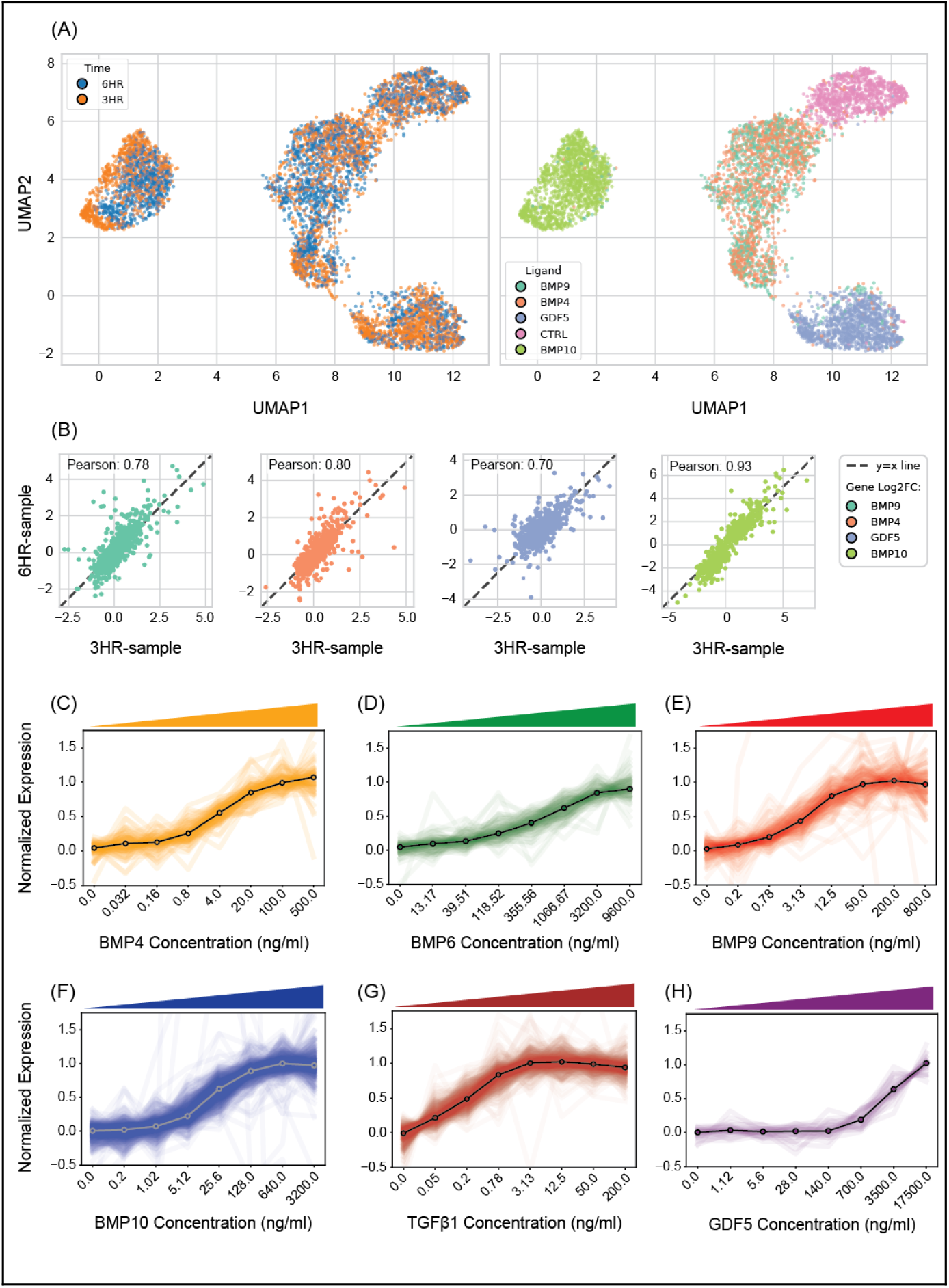
Temporal robustness and dose-dependent transcriptional responses across BMP/TGFβ ligands. (A) Uniform Manifold Approximation and Projection (UMAP) of single-cell transcriptomes from 3-hour and 6-hour stimulation experiments. Left: Cells colored by culture duration. Right: Cells colored by ligand treatment, illustrating the global separation of transcriptional states across signaling conditions. (B) Comparison of differential gene expression between 3h and 6h samples; each point represents a fold-change value of a responsive gene. The observed strong correlations indicate consistent transcriptional responses across timepoints. (C–H) Dose-response profiles for ligand-responsive gene sets. Lines show mean normalized expression of responsive genes for BMP4 (83 genes), BMP6 (101 genes), BMP9 (147 genes), BMP10 (614 genes), TGFβ-1 (501 genes), and GDF5 (27 genes). Panels display the expression trajectory across increasing ligand concentrations, with colors indicating ligands identity. Across ligands, responsive genes exhibit coordinated, graded responses that follow a shared sigmoidal trajectory, with differences primarily in response magnitude rather than response shape.

**Figure S2.**
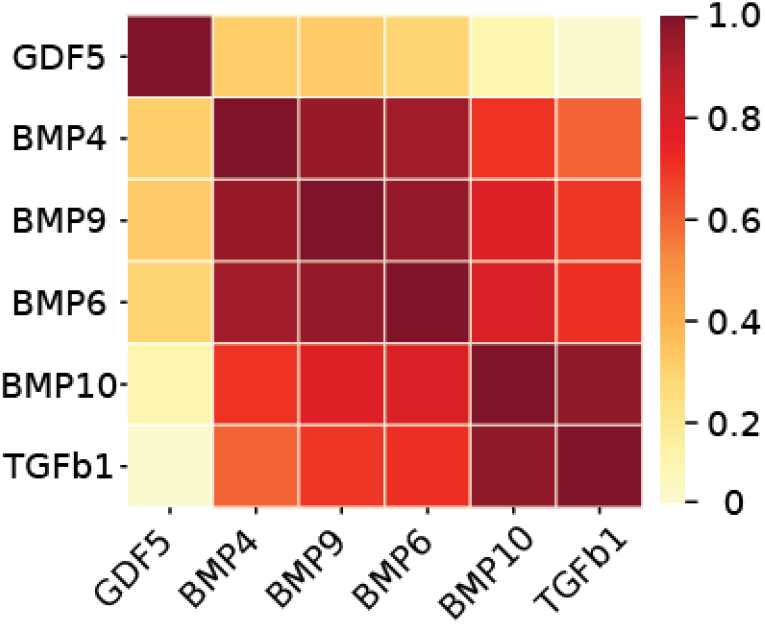
Correlation structure reveals grouping of BMP/TGFβ ligands into distinct transcriptional programs. Hierarchically clustered heatmap of pairwise Pearson correlations between ligands, computed from the log_2_ fold change values of all responsive genes (Figure 2B). The matrix reveals clustering of BMP4, BMP6, and BMP9 into a shared core BMP group, with BMP10 and TGFβ1 separating into an adjacent but distinct cluster, and GDF5 exhibiting a separate transcriptional profile.

**Figure S3.**
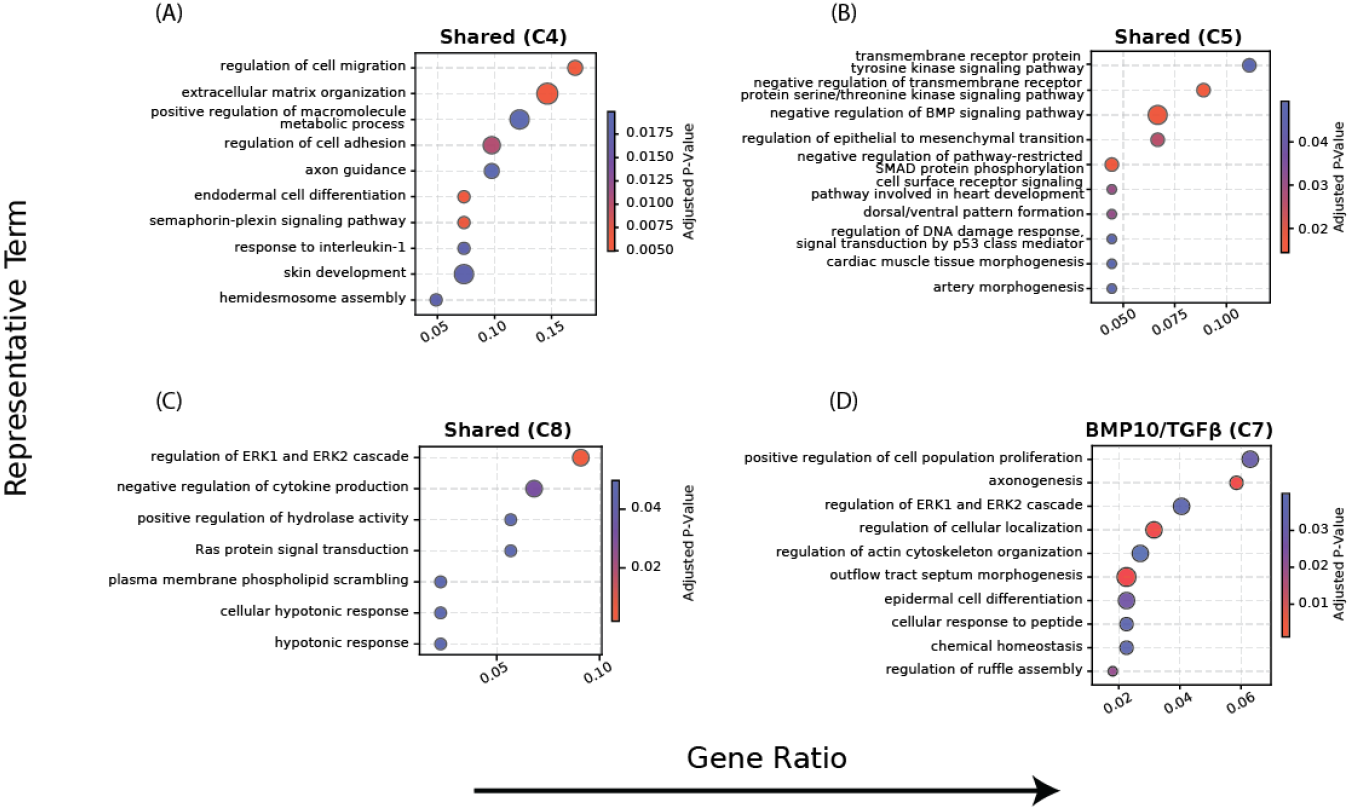
Gene Ontology analysis reveals distinct functional programs associated with shared and BMP10/TGFβ-responsive clusters. (A–D) GO enrichment analysis of clusters 6, 7, 8, and 9. Terms shown represent grouped GO categories, where representative terms were selected based on semantic similarity clustering (see Methods). Terms are ordered by gene ratio and colored by adjusted p-value.

**Figure S4.**
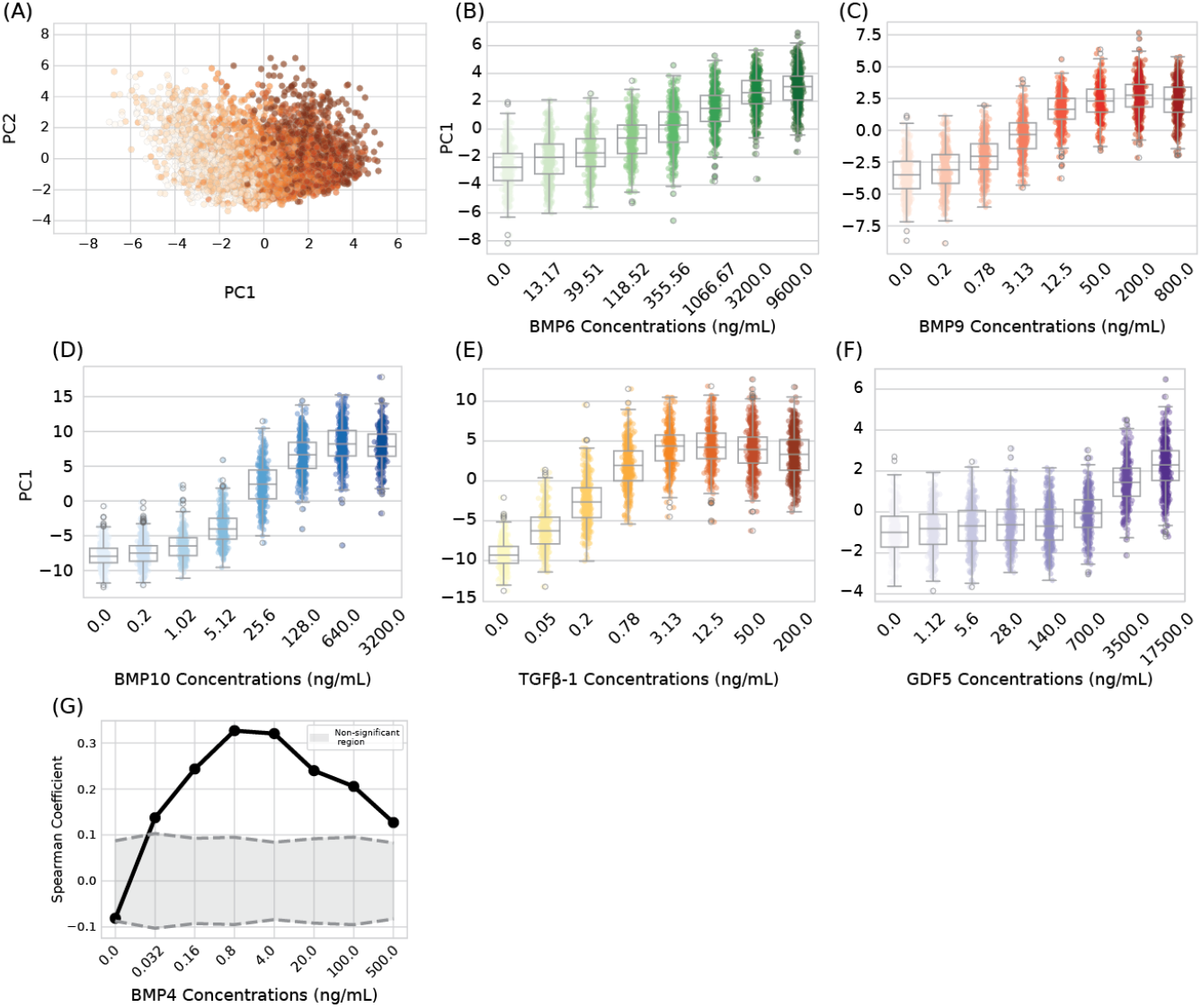
Perception score captures dose-dependent pathway activity across ligands. (A) PC1 and PC2 of BMP4 single cells projected on the mean response space of all concentrations, the shift in single cell state is visible across PC1. (B-F) Single-cell perception across the response range of all variants in our experiment. (G) Spearman correlation between perception score and BMP reporter within BMP4 concentrations, showing the highest correlation in the middle of the response range

**Figure S5.**
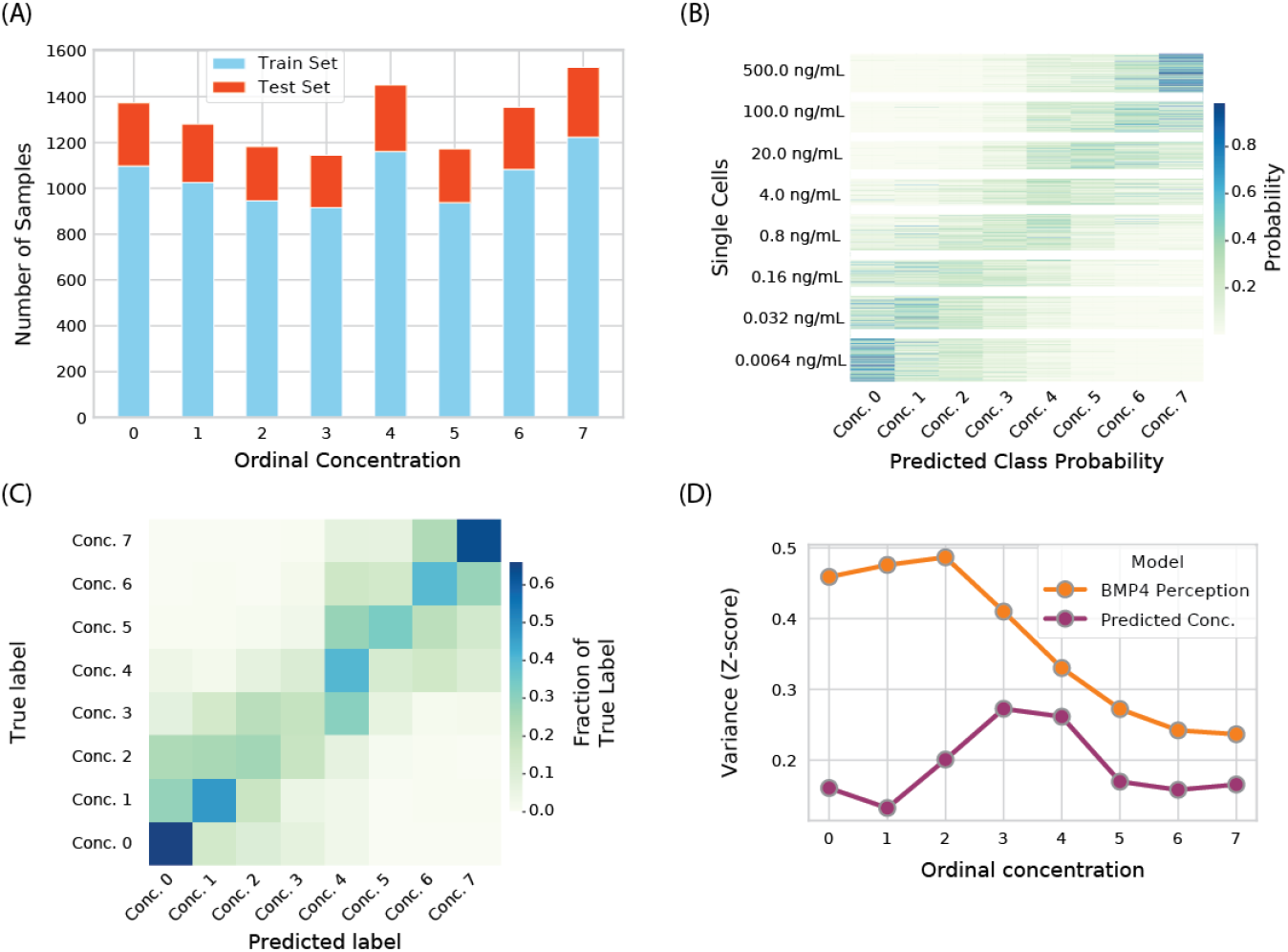
Concentration prediction using machine learning. (A) Split between train (light blue bars) and test (orange bars) sets across all ordinal classes in our data. (B) Prediction probabilities for each cell in the train set across all ordinal concentrations. Cells are grouped by their true class. (C) Confusion matrix for the XGBoost model was calculated for the test set. (D) Variance comparison of standardized values of BMP4 Perception score and Predicted BMP concentration. Machine learning models show lower variance across all classes, indicating their greater reliability.

**Table S1.** Ligands and concentrations.

| **Table S1. Ligands and concentrations** |  |  |  |  |  |
| --- | --- | --- | --- | --- | --- |
| BMP4<br>(ng/ml) | BMP6<br>(ng/ml) | BMP9<br>(ng/ml) | BMP10<br>(ng/ml) | GDF5<br>(ng/ml) | TGFβ-1<br>(ng/ml) |
| 0 | 0 | 0 | 0 | 0 | 0 |
| 0.032 | 13.17 | 0.2 | 0.2 | 1.12 | 0.05 |
| 0.16 | 39.51 | 0.78 | 1.02 | 5.6 | 0.2 |
| 0.8 | 118.52 | 3.13 | 5.12 | 28 | 0.78 |
| 4 | 355.56 | 12.5 | 25.6 | 140 | 3.13 |
| 20 | 1066.67 | 50 | 128 | 700 | 12.5 |
| 100 | 3200 | 200 | 640 | 3500 | 50 |
| 500 | 9600 | 800 | 3200 | 17500 | 200 |

**Table S2.** Specification of ligands used.

| **Table S2. Specification of ligands used.** |  |  |
| --- | --- | --- |
| **Ligand name** | **Vendor** | **Cat #** |
| BMP4 | R&D Systems | 5020-BP-010 |
| BMP6 | R&D Systems | 6325-BM-020 |
| BMP9 | R&D Systems | 5566-BP-010 |
| BMP10 | R&D Systems | 6038-BP-025 |
| GDF5 | R&D Systems | 853-G5-050 |
| TGFβ1 | R&D Systems | 7666-MB-005 |

